# Zero-Shot Generalization of Prospective Planning in Macaques

**DOI:** 10.64898/2026.10.19.760148

**Authors:** Myeong Seong Bak, Bowen Zhu, Yuki Minai, John J. Sakon, Gouki Okazawa, Roozbeh Kiani

## Abstract

Achieving a goal in natural environments often requires a sequence of actions whose consequences extend beyond their immediate outcomes. Because actions within a sequence can hinder or facilitate subsequent actions, successful behavior often demands prospective planning: selecting current actions based on their consequences for future actions. Although prospective planning is well documented in humans, its generalization to unfamiliar problems in nonhuman animals remains poorly understood. Here, we developed maze-navigation tasks in which macaque monkeys moved a token toward a goal while navigating around obstacles. Monkeys selected routes based on the positions of downstream obstacles, and their choices could not be explained by simple whole-maze visual summaries. In previously unseen, complex branching mazes, monkeys avoided deceptive dead ends—routes that initially approached the goal but ultimately failed—even when those dead ends extended several maze segments downstream. Prospective route selection was evident from the first encounters with multistep mazes and generalized to previously unencountered arrangements of familiar obstacles. These findings demonstrate zero-shot generalization of prospective planning: macaques applied a prospective strategy to unfamiliar multistep problems without learning the appropriate action sequences through repeated experience.

## Introduction

Achieving goals in natural environments often requires a sequence of interdependent actions. Each step may constrain or facilitate those that follow. As a result, optimal behavior demands *prospective planning*: selecting current actions based on their future consequences, not just immediate outcomes. For instance, a driver’s choice at the next intersection depends not only on the turn itself, but also on the downstream traffic flow and the route options it enables. Planning ahead—evaluating how actions unfold over time—is critical to reaching long-term goals efficiently.

In humans, prospective decision strategies have been studied across a range of settings, from elaborate laboratory tasks to complex games like chess (Campitelli and Gobet 2004; Chase and Simon 1973; Ho et al. 2022; Huys et al. 2012, 2015; Kolling et al. 2018; Miller and Venditto 2021; Reingold et al. 2001; Sheridan and Reingold 2017; Snider et al. 2015; Van Harreveld et al. 2007; Van Opheusden et al. 2023; van Opheusden et al. 2021). Multi-step decision tasks in the laboratory reveal whether people prioritize long-term value over short-term gain, and how deeply they plan into the future (Huys et al. 2012, 2015; Kolling et al. 2018; Snider et al. 2015; Van Opheusden et al. 2023; van Opheusden et al. 2021). Since the cost of immediate action could be superseded by the reward of future actions, these tasks can assess if humans find solutions that yield a better overall outcome through prospective planning. Similarly, strategic games with complex decision tree structures have been used to quantify human planning depth (Campitelli and Gobet 2004; Chase and Simon 1973; Reingold et al. 2001; Sheridan and Reingold 2017; Van Harreveld et al. 2007) and its refinement with training (Campitelli and Gobet 2004; van Opheusden et al. 2021). These same paradigms now serve as benchmarks for machine learning models (Schrittwieser et al. 2020; Silver et al. 2017, 2018), underscoring the computational richness of prospective planning and its necessity for intelligent behavior.

By contrast, prospective planning in nonhuman animals remains poorly understood, partly due to the challenge of adopting similarly complex tasks for them. Monkeys can learn action sequences, such as saccade chains or movement patterns (Averbeck and Lee 2007; Averbeck et al. 2006; Mushiake et al. 2001, 2006), and exhibit sophisticated hierarchical strategies in complex games (Yang et al. 2022). Other studies have examined model-based learning in sequential decision tasks with fixed state transitions (Miranda et al. 2020) and revealed rapid learning and efficient exploration in complex rodent mazes (Rosenberg et al. 2021). Together, these studies demonstrate remarkable flexibility in animal behavior, but leave open how readily animals can apply prospective strategies to unfamiliar multistep problems. Moreover, many paradigms require extensive training, making it difficult to distinguish the acquisition of task-specific strategies from the application of previously acquired cognitive abilities to new problems. An important open question is whether animals can exhibit zero-shot generalization of prospective planning: applying previously acquired knowledge of action consequences to unfamiliar multistep problems without learning configuration-specific solutions through repeated experience.

To address these limitations, we developed maze-navigation tasks that probe prospective planning in macaque monkeys using naturalistic interactions with a touchscreen and minimal task-specific training. In the main task, monkeys moved a token toward a goal while navigating around three immobile obstacles (walls) whose positions varied randomly across trials. The route chosen around one obstacle could hinder or facilitate subsequent navigation, allowing us to test whether current choices depended on future obstacles. We found that monkeys selected routes based on the sequential arrangement of downstream walls, rather than solely on global visual properties of the maze. In a second task involving more complex, branching mazes, monkeys avoided dead ends even when those routes offered greater immediate progress toward the goal. Critically, prospective route selection was evident from the monkeys’ first encounters with multistep mazes and generalized to previously unencountered configurations of familiar obstacles. Our findings suggest that monkeys can transfer previously acquired cognitive abilities to novel multistep problems, providing evidence for zero-shot generalization of prospective planning.

## Results

### Monkeys plan prospectively in a maze navigation task

We developed two maze-navigation tasks to test whether monkeys select current routes based on their consequences for later navigation. We use prospective planning to describe this sensitivity to downstream consequences, without assuming a particular mechanism for evaluating future paths. Both tasks were performed on a touchscreen installed on each monkey’s home cage (Fig. 1A). In the main task, monkeys swiped to move a token (a banana icon) from a starting position on one side of the screen to a green goal region on the opposite side while navigating around three immobile obstacles (walls) positioned randomly between the starting point and goal (Fig. 1B). To pass each wall, monkeys steered the token through either the upper or lower gap between the wall and the maze boundary, provided that the gap was sufficiently large. This task allowed us to test prospective route selection. In the example shown in Fig. 1C, the first wall could be passed from either above or below. However, because the second wall blocks the upper route, passing the first wall from above requires a sharp downward turn that elongates the total path (dashed red arrow). Passing the first wall from below instead allows the token to continue along the lower track with fewer directional changes and a shorter travel distance (solid red arrow). More generally, the route around a wall that minimizes subsequent track switches depends not only on the current wall but also on downstream wall positions.

We trained three monkeys to perform the maze navigation task. During initial training, monkeys encountered a single wall and were rewarded for moving the token to the goal. This phase taught the monkey the task logic—the properties of the token, goal, wall, and basic action-outcome mapping—without requiring them to incorporate downstream obstacles into their route choice. We then introduced multi-wall mazes and tracked behavior across thousands of trials. In this section, we analyze the final 20% of three-wall trials, a period after extensive task exposure when changes in strategy were unlikely (19,330 trials: monkey L, 7,002; M, 6,625; H, 5,703). Monkeys completed nearly all of these trials successfully (monkey L, 99.3%; M, 97.8%; H, 99.5%), with average completion times below 2 s (monkey L, 1.01 s; M, 1.67 s; H, 1.44 s). Subsequent sections will examine behavior immediately after multi-wall mazes were introduced and its evolution with continued task experience.

Choices around the first wall depended on the positions of all three walls (Fig. 1D; positions normalized to range between –0.5 and +0.5, with positive values indicating larger upper gaps; see Fig. S1A for individual monkeys). Larger upper gaps at the first wall increased the probability of choosing the upper route. Crucially, larger upper gaps at the second and third walls also increased the probability of passing the first wall from above. We quantified these effects using a logistic regression in which the probability of choosing the upper route around the first wall was modeled as a function of all three wall positions (Eq. 1). The model captured behavior well (Fig. 1D; *R*^2^ = 0.985), and revealed statistically significant effects for all three walls, with stronger effects for the more proximal walls (*β*_11_ = 11.325 *±* 0.169; *β*_21_ = 5.601 *±* 0.117; *β*_31_ = 2.358 *±*0.096; Wald test *p <* 10^−10^ for all). Individual monkeys showed the same pattern (Fig. S2A), indicating that each monkey incorporated information about future obstacles into its initial route choice.

Because the token could pass any gap larger than its size, we asked whether wall position had a saturating effect on choice. We therefore fit an alternative model in which wall positions passed through a compressive transformation before entering the logistic regression (a sigmoid transformation was used to compress larger gaps on both upper and lower sides; Eq. 10). The fitted transformation showed only modest compression at the largest gap sizes (Fig. S2B), and the linear and non-linear models generated nearly identical predictions for the monkey’s behavior (Fig. S2C). Thus, over the tested range, the influence of gap size was approximately linear.

We next asked whether the monkey’s forward-looking choices reduced subsequent track switching. We quantified track switching by the number of switches between the upper and lower tracks, a measure that was directly related to path length (Fig. 1B). To quantify the number of switches on each trial, we classified each wall position as U if only its upper gap was passable, L if only its lower gap was passable, or B if both gaps were passable. Each maze could therefore be described by a three-letter code, such as BLB in Fig. 1C (letters are ordered from wall 1 to 3). We then defined the lowest-switch route for each configuration. In the BLB example, this route passes the first wall from below (solid red arrow in Fig. 1C).

Monkeys chose the lowest-switch route on more than 80% of trials (monkey L, 81.0%; M, 94.0%; H, 86.0%; z-test against 50% chance level, *p <* 10^−10^ for all monkeys; only mazes with a unique lowest-switch route were included). Performance depended on maze configuration (Fig. 1E for monkey L, and Fig. S3A for monkeys M and H). For BUU and BLL mazes, where walls 2 and 3 favored the same route, monkeys were nearly perfect (BUU, 93.9%; BLL, 87.4%). For BUL and BLU mazes, which required a track switch, performance was lower (BUL, 84.4%; BLU, 67.4%). Part of this variation reflected a small but systematic bias toward upper routes, captured by a significantly positive intercept term in the logistic model (*β*_01_ = 0.82 *±* 0.028, *p <* 10^−10^). This bias was present in each monkey (Fig. S3A) and may have arisen from the position of the touchscreen relative to the elbow in our setup, which facilitated upward swipes. Nevertheless, downstream wall positions strongly modulated choices over and above this directional bias.

Choices around the second wall also depended on the position of the third wall and on the route taken around the first wall. After passing the first wall along one track, monkeys tended to remain on that track when possible. We extended the logistic model to include both the preceding route and downstream wall positions (Eqs. 7, 8; Fig. 2B,D). The preceding route had a strong influence on subsequent choices (wall 2, *β*_12_ = 11.95 *±* 0.27; wall 3, *β*_23_ = 10.35 *±* 0.22; *p <* 10^−10^ for both). Critically, the position of wall 3 also influenced the route chosen around wall 2 (*β*_32_ = 1.54 *±* 0.14, *p <* 10^−10^; Fig. 2B). These choices reduced unnecessary track switches and frequently followed the minimum-switch route (Fig. 2C, E, Fig. S3B, C).

**Figure 1.**
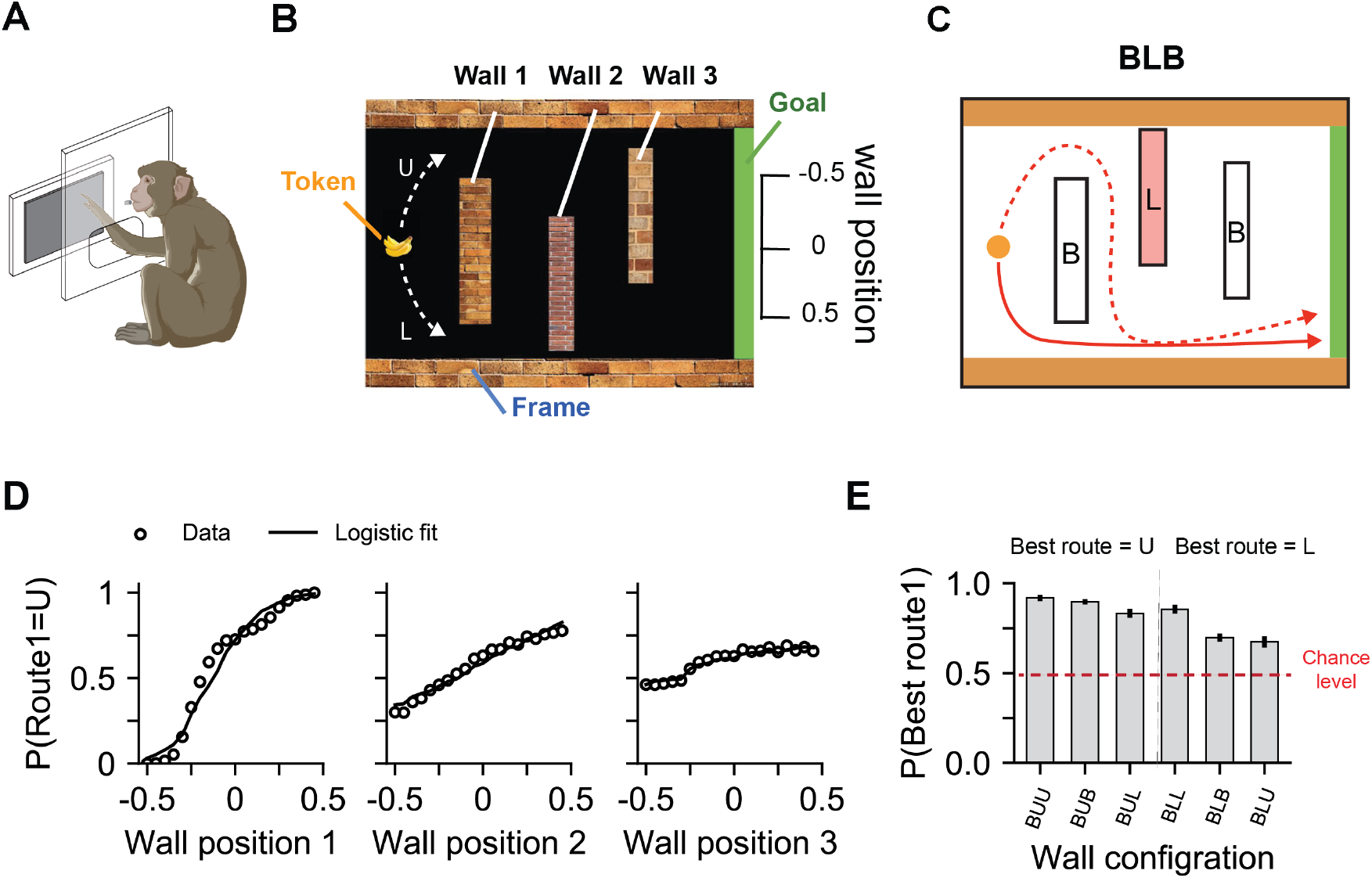
Prospective planning in a maze navigation task. (A) Experimental setup. Monkeys performed a maze navigation task using a touchscreen attached to the home cage. Juice rewards were delivered via a spout while the monkey performed the task (Illustration created with BioRender.com.) **(B)** Task display. On each trial, monkeys moved a token (banana icon) from one side of the screen to a green goal region on the opposite side to obtain reward. The token could not pass through the maze boundaries or the three immobile obstacle walls. **(C)** Task structure requires prospective planning. The route chosen around one wall affected the ease of passing subsequent walls. Each wall’s position was categorized based on the passable gaps around it: U (upper only), L (lower only), or B (both). Each maze configuration could then be defined using a three-letter code (e.g., BLB). **(D)** Monkeys’ decisions were influenced by future wall positions. The probability of choosing the upper route around the first wall (Route1) is plotted as a function of the three wall positions (open circles; wall center positions normalized to [-0.5, 0.5] and binned into 20 quantiles; positive positions indicate larger upper gaps). Solid lines show the fit of a logistic regression model (Eq. 1). See Fig. S1A for individual monkeys’ results. **(E)** Monkeys favored lowest-switch paths. They preferentially chose routes that minimized sharp turns and total path length. Data are shown for monkey L. All monkeys showed qualitatively similar behavior (Fig. S3A). The red line indicates chance level (50%). Error bars denote SEM.

**Figure 2.**
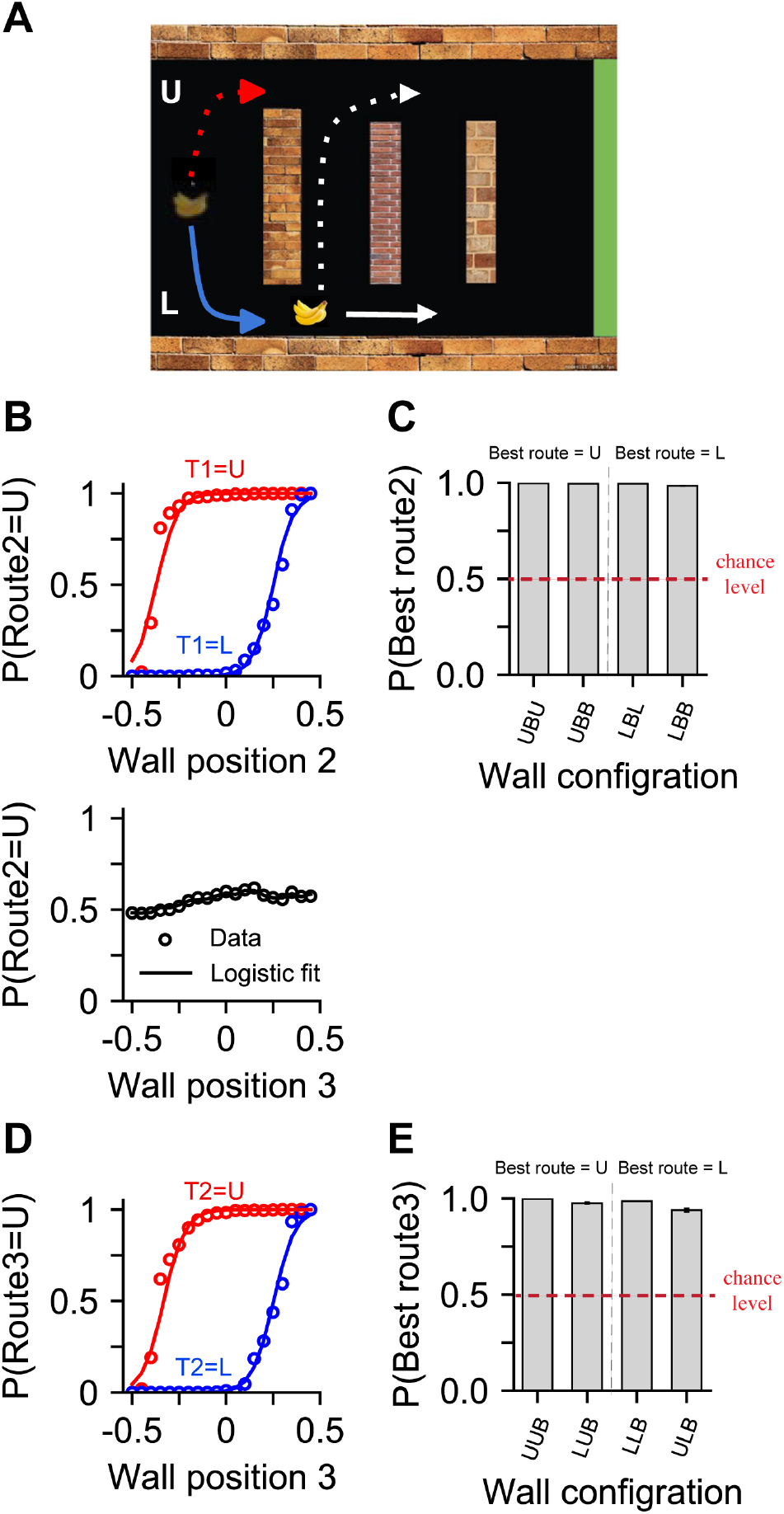
Selected routes around the second and third walls depended on the chosen route for the preceding wall. (A) Sequential path constraints. The route taken around a prior wall (e.g., the lower route at wall 1) determines the token’s position before approaching the next wall. We tested how monkeys’ choices around walls 2 and 3 were influenced by the current wall’s position and the preceding route. Some paths avoided sharp turns and reduced total travel distance (e.g., solid white arrow vs. dashed arrow). **(B)** Prior choices and future wall positions influenced choices at the second wall. The probability of choosing the upper route at wall 2 varied with the monkey’s route at wall 1 (T_1_) and the positions of walls 2 and 3. Data are aggregated across all monkeys. **(C)** Monkeys selected lowest-switch routes around wall 2, for nearly all wall configurations. Our analysis focuses on maze configurations in which wall 2 was B, allowing the monkey to successfully complete the maze using either U or L routes around wall 2. Data shown for monkey L. Similar patterns were observed in the other monkeys (Fig. S3B). The first letter (U or L) in each three-letter code indicates the route taken at wall 1, rather than the wall position. Error bars represent SEM. **(D)** Choices at the third wall were influenced by wall 3 position and the route taken at wall 2 (T_2_). **(E)** Monkeys chose lowest-switch routes at wall 3. The first two letters in the three-letter codes are based on the actual routes taken at walls 1 and 2. See Fig. S1B-C and S3B-C for individual monkeys’ results.

### Wall-specific prospective choices cannot be explained by whole-maze visual heuristics

Because the entire maze was visible at the start of each trial, the influence of downstream walls on the monkey’s initial route choice does not by itself establish wall-specific prospective planning. An alternative possibility is that monkeys reduced the complete maze to a single visual summary—for example, whether the upper or lower half of the display appeared more open—and selected the initial route accordingly. Such a strategy would also make choices around the first wall depend on walls 2 and 3, without requiring the monkey to preserve the identity or sequential position of individual obstacles.

We tested this possibility by comparing the wall-specific model (Eq. 1) with two models that represented the maze configuration using order-free summaries of upper and lower lane openness. These alternatives included a model that used the average vertical position of the three walls (average-position model; Eq. 3) and a model that used the largest upper gap minus the largest lower gap across the three walls (largest-gap model; Eq. 5). Unlike the wall-specific model, these summary models discarded the identity and serial position of individual walls, representing the maze through aggregate visual statistics. We compared the models using leave-one-session-out cross-validation performed separately for each monkey. For each iteration, the models were fitted to all but one data-collection session and evaluated on the held-out session, repeating this procedure until every session had been tested once. The wall-specific model predicted the monkeys’ choices around the first wall substantially better than either summary model (cross-validated *R*^2^, wall-specific model versus the better-performing summary model: H, 0.691 versus 0.413; L, 0.590 versus 0.324; M, 0.566 versus 0.419).

The difference was especially pronounced for maze configurations in which the same aggregate measures of openness arose from different arrangements of individual walls. Specifically, we examined BUL and BLU configurations, which had the same wall types, average wall position, and overall amounts of obstruction in the upper and lower portions of the maze, but imposed different sequential constraints. Thus, they provided a direct test of whether route choices depended on the order of individual wall positions rather than the aggregate visual statistics captured by the summary models. We focused on monkey L because it was the only monkey with both configurations in the analyzed main-task data; BLU was reserved for generalization testing in H and M. Monkey L preferred the upper route in BUL mazes (83.3% of 228 trials) and the lower route in BLU mazes (67.4% of 230 trials). The wall-specific model predicted these choices better than either summary model (cross-validated *R*^2^ = 0.394, compared with 0.102 for the better-performing summary model).

A prospective account predicts that the monkey evaluates the route forward: to determine whether to pass above wall 1, it must consider the route options available at wall 2 and how those options shape the relevance of wall 3. Having established that all three walls contributed to the initial route choice (Eq. 1), we next asked whether the influence of wall 3 depended on the routes permitted by wall 2. Specifically, when wall 2 permits only one route (upper or lower), it already constrains the token’s subsequent trajectory, potentially reducing the influence of wall 3 on the initial choice compared with configurations in which wall 2 permits both routes. To test this prediction, we extended Eq. 1 with an interaction between wall 3 position and whether wall 2 permitted one or both routes (Eq. 2). Adding this interaction improved model fit (likelihood-ratio test, *χ*^2^(1) = 69.55, *p* = 7.5 *×* 10^−17^). The interaction coefficient was negative (*β*_41_ = −1.49 *±* 0.18), indicating that restricting passage at wall 2 reduced the influence of wall 3 on the initial choice by ∼50% (relative to its logistic regression coefficient when wall 2 permitted both routes). The interaction coefficient was negative in all three monkeys and significant in two (*p <* 0.001). Thus, the influence of wall 3 on the initial choice depended on the routes available at the intervening wall. This finding is consistent with monkeys considering how successive obstacles constrain future actions.

Together, these results argue against strategies based solely on a global, Gestalt-like estimate of which side of the maze appeared more open. The monkeys’ initial choices depended on the identity and order of down-stream walls, and the influence of individual walls varied with the route constraints imposed by intervening obstacles. These findings are consistent with prospective evaluation of how current choices constrain subsequent navigation. However, we note that these analyses do not identify the particular computations used by the monkeys. Rather, they establish an important property of the monkeys’ strategy: current choices reflected the sequential arrangement of future obstacles and could not be explained by the order-free whole-maze visual summaries tested here.

### Prospective route selection in multicursal mazes

To test prospective route selection in a richer setting, two monkeys (M and H) also performed a path-finding task with substantially more complex mazes. On each trial, the monkey was shown a multicursal maze and was rewarded for successfully dragging a token—a raisin icon—to a green goal region. The token could not cross the maze walls. Mazes were generated anew on each trial, and rare repetitions were excluded from analysis, precluding monkeys from relying on repeated exposure and memorization of particular displays or action sequences. The mazes contained multiple branching paths and numerous dead ends (e.g., Fig. 3A). At each branch, the monkey therefore had to choose between paths with different downstream consequences. Entering a dead end required reversing course and retracing part of the route, making exploratory selection costly in time and movement. Monkeys were initially familiarized with simple 2 *×* 2 mazes, where prospective planning was not required, and were subsequently tested with more complex 6 *×* 6 and 7 *×* 7 mazes.

Monkeys successfully completed 65,518 of 70,343 structurally unique mazes (93.1%; H, 92.8%; M, 93.5%; session-cluster bootstrap 95% CI, 92.4–93.8%; Fig. 3B). A comprehensive characterization of this dataset lies beyond the scope of the present study. Here, we asked whether monkeys selected routes that avoided dead ends. We examined branching points at which the monkey could choose between a route that eventually reached the goal and one or more routes that terminated in dead ends. Because the mazes contained no loops, only one forward route at each such branching point could reach the goal without retracing the path. We analyzed the monkey’s first encounter with each branching point, excluding backtracking to the preceding maze cell. Across 307,874 such encounters, monkeys chose the goal-reaching route on 81.8% of choices, compared with a mean random-choice level of 45.9% (difference, 35.9%; session-cluster bootstrap 95% CI, 35.4–36.4%; one-sided node-specific *Z*-test, *z* = 403.77, *p <* 10^−12^; H, 82.4%; M, 81.3%; Fig. 3C).

We next examined whether monkeys could avoid dead ends when doing so required rejecting the route most directly oriented toward the goal. A simple goal-direction heuristic would select, at each branching point, the available step that pointed most directly toward the goal, without considering where the path eventually led. Such a strategy would fail when the most directly goal-oriented step entered a dead end. To identify those cases, we measured the angle between each available forward (non-backtracking) step and the straight line connecting the branching point to the goal (Fig. 3D). The step forming the smallest angle was most directly oriented toward the goal. We defined a deceptive dead end as a dead-end branch whose initial step formed a uniquely smaller angle than the other available forward steps. Avoiding a deceptive dead end therefore required rejecting the option favored by the local heuristic in favor of a route that could ultimately reach the goal. We further asked whether monkeys avoided deceptive dead ends when their adverse consequences lay farther downstream. We quantified dead-end depth as the number of maze-cell transitions from the branching point to the farthest terminal cell within the deceptive dead-end branch, including any subbranches. Shallow dead ends terminated near the current choice point, whereas deeper dead ends extended farther downstream. Thus, dead-end depth provided a measure of the spatial extent of the path structure relevant to evaluating a deceptive dead end.

Among 78,407 first encounters with branching points containing a deceptive dead end, monkeys chose the goal-reaching route above the node-specific random-choice level at every dead-end depth (Fig. 3E; depth 1, 71.1% versus 44.4%, *n* = 34,387; depth 2, 68.9% versus 44.5%, *n* = 13,950; depth 3, 68.5% versus 44.5%, *n* = 7,382; depth *≥* 4, 66.3% versus 44.0%, *n* = 22,688; one-sided node-specific *z*-tests, all *z ≥* 42.11, all *p <* 10^−300^; session-cluster bootstrap 95% CIs for the accuracy-minus-chance difference excluded zero at every depth). Thus, monkeys did not simply select the route that produced the greatest immediate progress toward the visible goal. Instead, they sacrificed short-term gains when necessary and used downstream path consequences to select actions that ultimately completed the maze, even when the rejected dead-end branches extended multiple maze cells beyond the current choice point. Importantly, these prospective choices occurred in structurally unique mazes that the monkeys had never encountered before. Thus, monkeys generalized their prospective route-selection strategy to novel arrangements of branching paths and dead ends without learning solutions to individual mazes through repeated experience.

**Figure 3.**
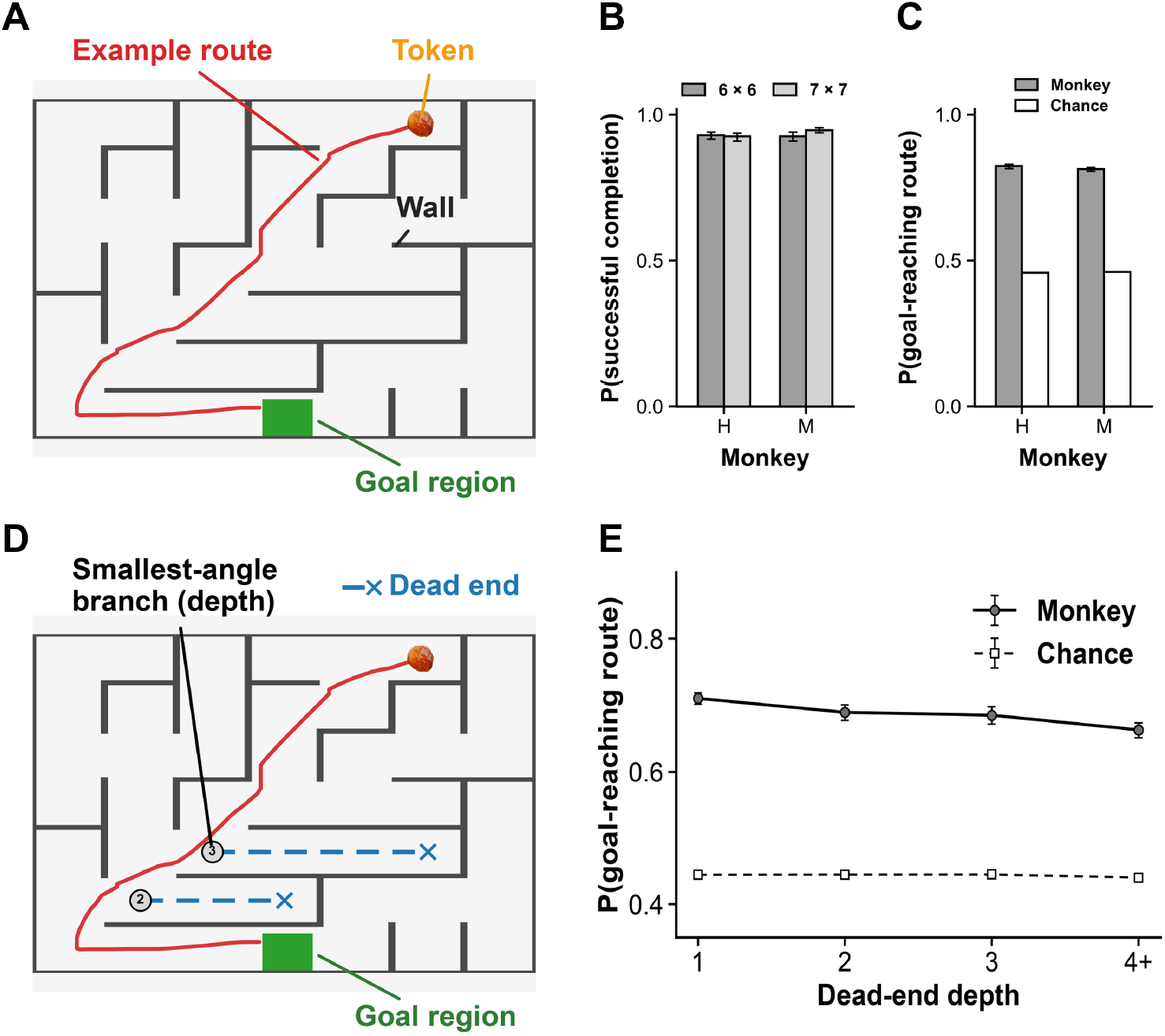
Monkeys selected goal-reaching routes in multicursal mazes. (A) Example maze showing the trajectory (red) the monkey used to move the token to the goal (green). Maze walls are not passable. **(B)** Completion probability for 6 × 6 and 7 × 7 mazes in monkeys H and M. **(C)** Probability of selecting the goal-reaching route at the maze branching points that offered both goal-reaching and dead-end forward options. **(D)** Definition of deceptive dead ends and dead-end depth. A local heuristic selects the available forward step forming the smallest angle with the direction of the goal. A deceptive dead end is a branch favored by this heuristic that ultimately terminates without reaching the goal. Dashed blue lines indicate deceptive dead-end branches, crosses mark their terminal cells, and circled numbers indicate dead-end depth (maximum number of cell transitions to a terminal cell). **(E)** Probability of selecting a goal-reaching route over a deceptive dead end, grouped by dead-end depth (4+: four or more cell transitions). Data are pooled across monkeys H and M. Chance is the mean of 1/k, where k is the number of available non-backtracking options, calculated separately for each monkey (C) or depth group (E). Error bars indicate session-cluster bootstrap 95% confidence intervals.

### Prospective planning without training: experience refined speed, not strategy

In both maze tasks, monkeys exhibited prospective route selection without repeated training on individual maze configurations. We examined how rapidly this strategy appeared in the three-wall task, for which behavior was recorded from the very first encounters with multiple walls (Fig. 4). Before these encounters, monkeys had been trained on single-wall mazes, which taught the visuomotor rules of the task but did not require choosing a current route based on downstream obstacles. Thus, prospective choices during initial multi-wall trials could not have been acquired through repeated experience with those particular configurations. Instead, monkeys appeared to apply previously acquired abilities to the novel problem of navigating multiple interacting obstacles.

Indeed, the monkey’s initial encounter with two- and three-wall mazes already bore the hallmarks of prospective planning. We analyzed the first choice around wall 1 in BX configurations for two-wall mazes and BXX configurations for three-wall mazes, where X denotes any wall type. In these trials, both routes were available around wall 1, and the route producing fewer subsequent switches depended on the downstream walls. On their *very first* encounter with each maze configuration, monkeys dominantly selected the lower-switch route (77% *±* 9%; p=0.008, binomial test against 0.5 chance level). Thus, prospective route selection was evident before repeated exposure to the tested multi-wall configurations.

Moreover, the selected route around wall 1 was modulated by all three wall positions in the first and final sessions (Fig. 4A). The principal wall-position effects were present in each monkey from the outset and showed no consistent increase over time (all *p >* 0.05, *z*-test for the difference in proportions, Bonferroni corrected). These results indicate that the structure of the planning strategy was already present when multi-wall testing began. The logistic regression parameters linking wall positions to route choices also remained largely stable across the experiment (Fig. 4B; Eqs. 1, 7, 8). Similarly, the likelihood of selecting the lower-switch route showed no systematic improvement with experience (Fig. 4C). Regression coefficients for the effect of experience were small and non-significant in each monkey (Eq. 11; Monkey M, *β*_*br*_ = 0.172 *±* 0.357, *p* = 0.63; Monkey L, *β*_*br*_ = 0.079 *±* 0.265, *p* = 0.77; Monkey H, *β*_*br*_ = 0.013 *±* 0.305, *p* = 0.97).

Experience nevertheless improved motor execution. Across sessions, monkeys completed mazes faster (Fig. 4D; Eq. 12; Monkey L, *β*_*st*_ = −0.035 *±* 0.005, *p* = 1.6 *×* 10^−4^; Monkey M, *β*_*st*_ = −0.064 *±* 0.004, *p* = 4.7 *×* 10^−7^; Monkey H, *β*_*st*_ = −0.060 *±* 0.007, *p* = 2.3 *×* 10^−5^). This speed-up originated from higher swiping speeds (Fig. 4E; Eq. 13; Monkey L, *β*_*ss*_ = 0.049 *±* 0.003, *p* = 2.6 *×* 10^−7^; Monkey M, *β*_*ss*_ = 0.047 *±* 0.006, *p* = 6.2 *×* 10^−5^; Monkey H, *β*_*ss*_ = 0.021 *±* 0.008, *p* = 0.023) and higher continuity of swipes, quantified by a decrease in the total number of swipes for reaching the goal in a trial (Fig. 4F; Eq. 14; Monkey L, *β*_*sc*_ = −0.047 *±*0.006, *p* = 7.7 *×*10^−5^; Monkey M, *β*_*sc*_ = −0.061 *±*0.003, *p* = 4.6 *×*10^−8^; Monkey H, *β*_*sc*_ = −0.059 *±* 0.008, *p* = 7.5 *×* 10^−5^).

Together, these results indicate that task experience refined motor execution but was not necessary for the emergence of prospective route selection in tested mazes. The monkeys appeared to draw on previously acquired cognitive abilities rather than learning a new planning strategy through repeated exposure to multistep mazes.

**Figure 4.**
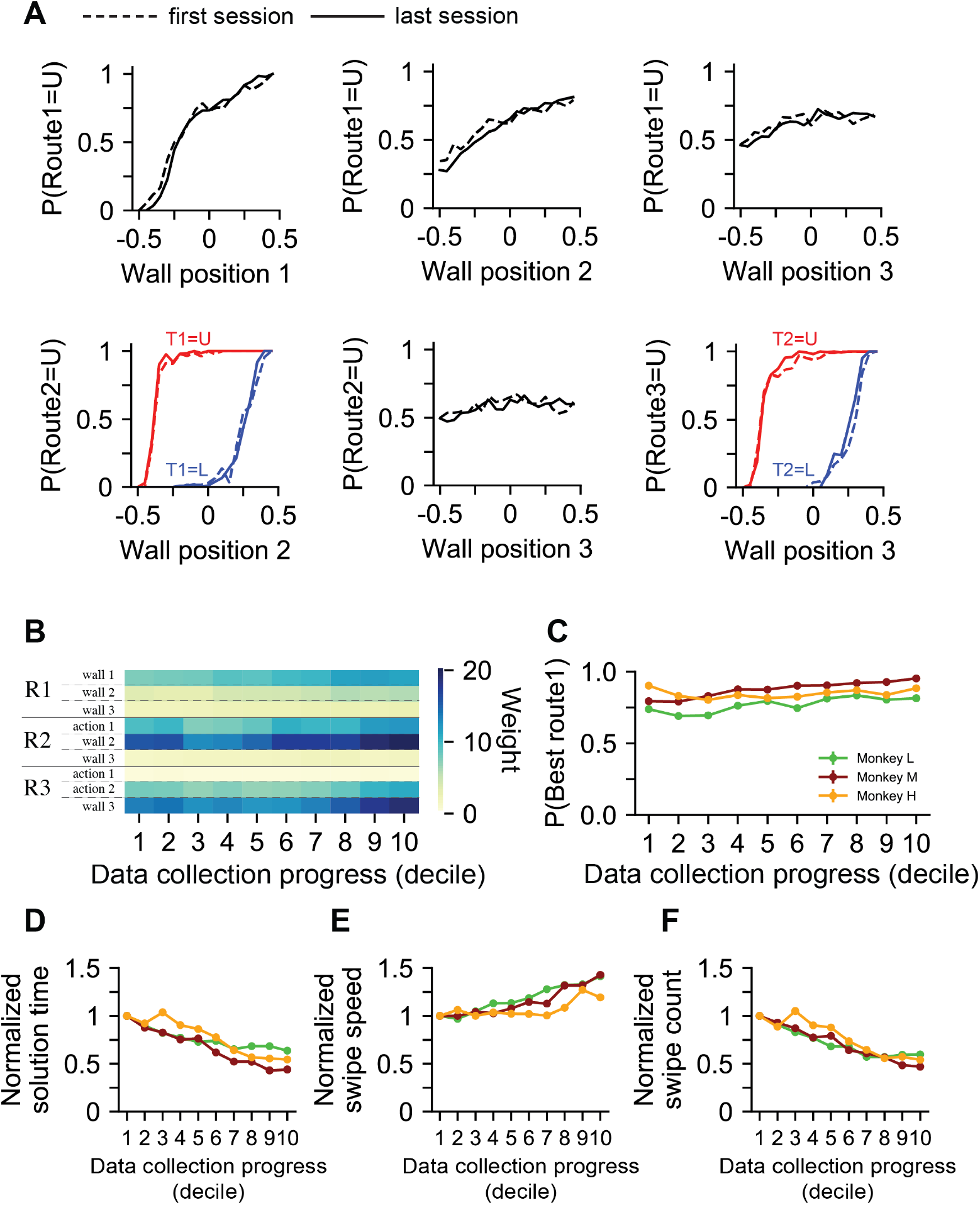
The prospective planning strategy was evident from the beginning of data collection. (A) Stability of strategy from the outset. Monkeys’ prospective planning strategy was present in the first session and remained stable through the last session. Results are aggregated across all three monkeys. **(B)** Model parameters remained stable over time. We divided the data into deciles based on collection order and fit logistic regression models within each decile (Eqs. 1, 7, and 8). Regression weights (colormap) showed minimal change over time. R1-R3 denote the three regression models for choices around walls 1-3. **(C)** No systematic improvement in lowest-switch route selection over the course of the experiment. **(D-F)** Experience improved motor execution. While planning strategy remained stable, motor performance improved with experience. Monkeys showed faster maze completion times (solution time; D), driven by increased swiping speed (E) and fewer swipes per trial (F). All values are normalized to the means for the first decile. Error bars denote SEM.

### Monkeys generalized their planning strategy to novel environments

The presence of prospective planning from the first multi-wall trials suggested that monkeys could generalize an existing prospective planning ability to the maze task. To further test their capacity for generalization, we conducted two control experiments with monkeys M and H. The controls examined generalization across changes that were either irrelevant or structurally relevant to path planning.

In the first control experiment, we altered a nuisance, task-irrelevant visual feature: the texture of the walls. During the main task, the three walls had distinct and fixed textures. During the control experiment, we randomized texture assignments across walls (Fig. 5A). If monkeys had memorized associations between complete visual displays and particular actions, changing the texture order could disrupt those associations. Instead, the monkey’s performance remained stable, with the probability of selecting the lower-switch route comparable before and after texture randomization (Fig. 5B). Thus, route selection was largely invariant to changes in task-irrelevant visual features, consistent with reliance on the structural properties of the maze.

In the second control experiment, we tested generalization to novel maze configurations. Of the 27 possible wall configurations (3^3^), only 19 were used regularly during the main data-collection sessions for monkeys M and H. The remaining eight were withheld from the main task for monkey H and presented only rarely to monkey M (1.7% of main-task trials). We tested generalization to these configurations during subsequent control sessions (Fig. 5C). Successful generalization would produce choices in these configurations that were predictable from the monkeys’ established sensitivity to individual wall positions.

**Figure 5.**
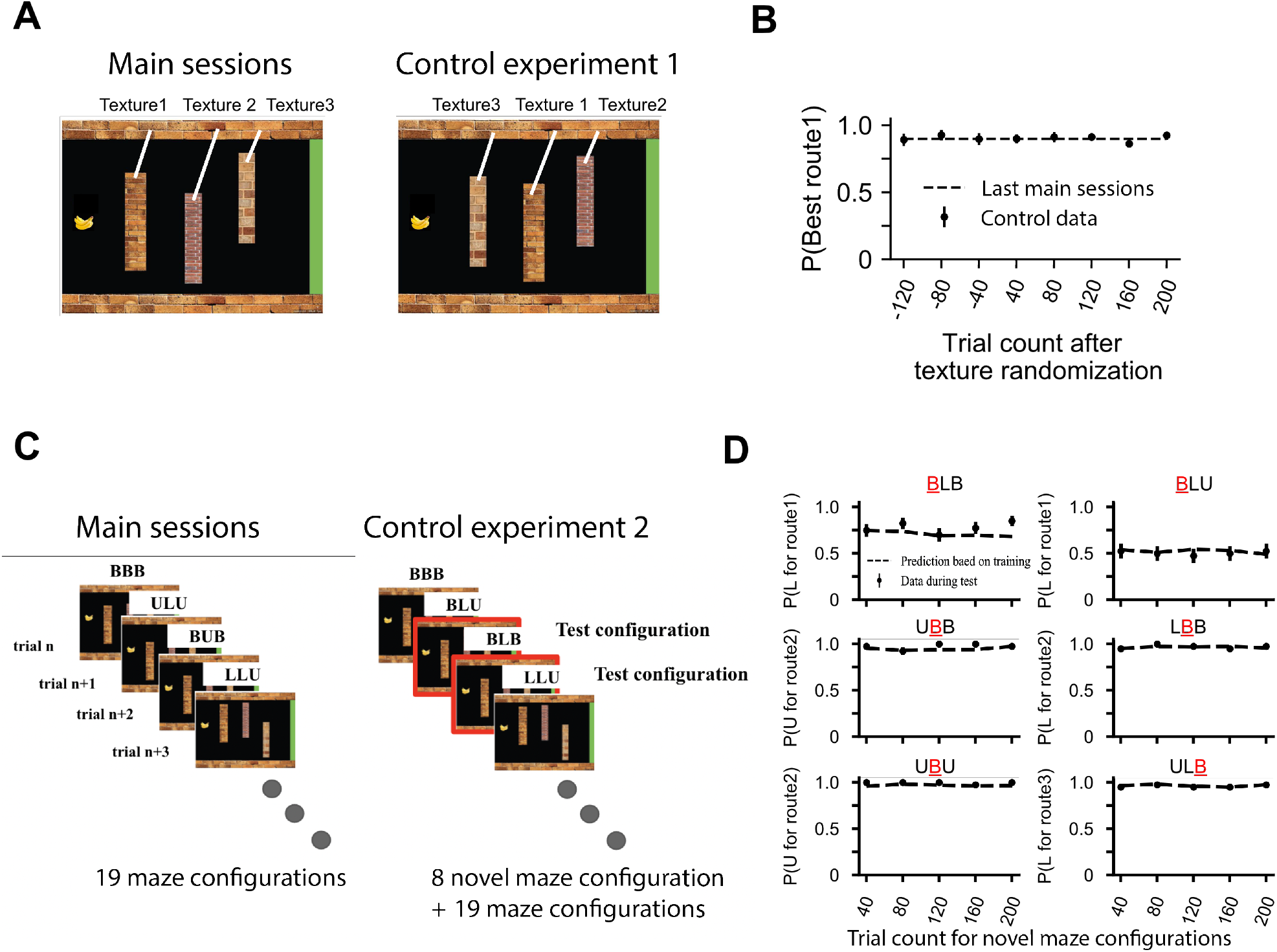
Monkeys generalized their strategy to novel conditions. (A) To examine whether monkeys generalized their planning strategy across visual display changes, we randomized the order of wall textures. In the main task, texture order was fixed. If monkeys had relied on specific associations between visual patterns and the best motor responses their performance would have deteriorated with texture randomization. **(B)** Monkeys’ performance remained unaltered despite the randomized wall textures. We binned trials before and after texture randomization (40 trials/bin) and plotted the probability of choosing the lowest-switch route around the first wall (data points). The dashed line represents the average performance in the last five sessions of the main experiment. **(C)** To examine generalization to structurally novel environments, we withheld 8 of the 27 possible maze configurations during the main task and introduced them only during a second control experiment (monkeys M and H). **(D)** Behavior for the held-out maze configurations closely matched predictions from a logistic regression model fit to the main task data. We binned trials involving the B-type walls in the control experiment (40 trials/bin; B-type walls permitted both U and L routes). Data points represent the observed choice probabilities. Dashed lines show model predictions based on observed behavior in the main task (Eqs. 1-8). Error bars denote SEM.

The monkey’s performance for the novel maze configurations was indeed comparable to that expected from their performance for the familiar configurations. We fit the logistic regression model to trials from the final sessions of the main task (Fig. 1, Fig. 2) and used them to predict choices in the novel configurations. For the six novel configurations containing at least one B wall, we compared predicted and observed choice probabilities (Fig. 5D). Behavior closely matched the out-of-sample predictions for the held-out configurations. This transfer is also consistent with the monkey’s performance in the multicursal maze task, where every test maze was unique.

Together, these results demonstrate that monkeys applied their prospective route selection strategy across changes in task-irrelevant visual features and to new combinations of familiar obstacle constraints. In particular, choices in configurations that were novel to monkey H and rarely encountered by monkey M could be predicted from their established sensitivity to individual wall positions, without repeated model training on those configurations. This provides evidence for zero-shot generalization of a prospective strategy, rather than memorization of specific displays or fixed movement sequences. The successful navigation of unique multicursal mazes further supports the flexibility of prospective route selection across unfamiliar spatial layouts.

## Discussion

Using two novel maze-navigation tasks, we show that macaque monkeys exhibit prospective planning: they consider downstream consequences of their choices for selecting current routes. In the three-wall task, monkeys’ initial choices depended on the positions and sequential arrangement of future obstacles, rather than simply on the overall visual openness of the maze. In more complex, branching mazes, monkeys avoided deceptive dead ends that initially directed the token toward the goal but ultimately prevented maze completion. These findings establish the prospective properties of the monkeys’ route-selection strategy. Critically, prospective route selection was evident from the monkeys’ first encounters with multistep mazes, and their strategy generalized to previously unencountered configurations. Together, these findings demonstrate that macaques can apply prospective planning to novel multistep problems without learning the appropriate action sequences through repeated experience.

Prospective planning has been extensively documented in humans, including in chess, multi-step decision tasks, and planning games, but evidence for similar capacities in nonhuman animals remains limited. Previous studies have shown that monkeys can learn sequences of actions, such as saccades or lever presses, and that neurons in prefrontal cortex can represent combinations of future actions (Averbeck and Lee 2007; Mushiake et al. 2006). Other studies have examined model-based learning in sequential decision tasks with relatively simple transition structures (Akam et al. 2021; Miller et al. 2017; Miranda et al. 2020). More complex behavioral paradigms have revealed hierarchical decision strategies in macaques (Yang et al. 2022) and and rapid learning in rodents navigating branching mazes (Rosenberg et al. 2021). Our experiments extend this work by examining whether monkeys can select routes based on downstream consequences in unfamiliar spatial environments. The ability to reject deceptive dead ends is particularly informative: monkeys sometimes sacrificed immediate progress toward the goal to avoid paths that could not lead to success, even when those dead ends extended several maze segments beyond the current choice point. These results provide behavioral evidence for prospective evaluation of alternative routes, although they do not establish whether monkeys explicitly simulate complete action sequences.

An important feature of our findings is the limited training required for prospective route selection. The monkeys learned the basic visuomotor rules of the task through interactions with the touchscreen and single-wall mazes. Nevertheless, their route choices already reflected downstream obstacles during their first encounters with multistep configurations. Moreover, experience improved movement speed and continuity without a corresponding systematic improvement in the structure of their route choices. These observations suggest that monkeys brought previously acquired cognitive abilities to the task rather than developing a new prospective strategy through repeated multistep training. This ability generalized to previously unencountered combinations of familiar wall constraints and, importantly, to structurally unique multicursal mazes containing numerous branching paths and dead ends. In the latter, monkeys routinely selected goal-reaching paths over deceptive dead ends despite having no prior experience with the particular maze layouts. Thus, the monkeys’ ability to use downstream consequences was not restricted to familiar configurations or to the relatively simple three-wall task. We interpret these findings as evidence for zero-shot generalization of prospective planning: the application of an existing planning capability to unfamiliar multistep environments without learning configuration-specific action sequences through repeated experience. This does not imply that the underlying cognitive abilities are innate or independent of prior experience.

What previously acquired abilities might support this transfer? Our paradigm engages a familiar form of interaction for macaque monkeys: manipulating objects with their hands to achieve a goal (Cabrera-Á lvarez and Clayton 2020). Navigating a token around obstacles may draw on cognitive abilities developed through extensive experience with physical objects and spatial relationships (Lake et al. 2017; Stachenfeld et al. 2017). These could include knowledge of movement constraints, collision avoidance, and the continuity of object trajectories. Consistent with the capacity of macaques to flexibly use structured spatial knowledge, recent work has demonstrated mental navigation between learned landmarks, including previously untested landmark pairs (Neupane et al. 2024). Such knowledge could help monkeys evaluate the consequences of alternative routes without learning each maze configuration independently. Our results do not establish the developmental origins of these abilities, but they suggest that naturalistic, spatially grounded tasks can reveal cognitive capacities that might be difficult to observe in paradigms requiring extensive training.

Although our task shares structural features with multi-step reinforcement learning paradigms (Akam et al. 2021; Dezfouli and Balleine 2019; Groman et al. 2019; Hasz and Redish 2018; Miller et al. 2017; Miranda et al. 2020), there are important differences. In our mazes, the complete spatial arrangement is visible before the monkey initiates navigation, allowing future path consequences to be evaluated without learning the transition structure of each configuration through trial and error. Moreover, choices are expressed through continuous, gesture-based movements rather than discrete responses between abstract states. The observed behavior is consistent with prospective evaluation of a spatial model of the environment, but the behavioral data do not uniquely identify the computations that implement this process. In particular, the monkeys might use explicit evaluation of alternative trajectories, a learned repertoire of visual-spatial rules, or other strategies sensitive to downstream path structure. Distinguishing these possibilities will require additional experiments and computational models.

One limitation of our study concerns the motor costs associated with different routes. In the three-wall task, we quantified trajectory complexity using the number of switches between upper and lower tracks, which was related to path length. However, fewer switches do not necessarily minimize the monkey’s subjective or energetic movement costs. Monkeys also exhibited a bias toward upper routes, potentially reflecting kinematic constraints imposed by the experimental setup. Future work could model movement costs more explicitly using trajectory kinematics. These considerations do not affect the central finding that choices depended on downstream obstacles, but they limit conclusions about whether monkeys selected globally optimal routes.

The neural and computational mechanisms of prospective planning remain open questions. One possibility is that monkeys rely on internal models that capture the physical constraints of the environment and allow prospective evaluation of possible trajectories (Battaglia et al. 2013; Smith and Vul 2013; Wolpert et al. 1995). Such models might draw on extensive developmental experience interacting with the physical world (Lake et al. 2017; Tenenbaum et al. 2011). Alternatively, prospective route selection could emerge from other computational strategies that preserve information about sequential path constraints. Although the descriptive models used here quantify which aspects of the maze influence choices, they do not distinguish these possibilities. Identifying the underlying computations and their neural substrates remains an important challenge (Miller and Venditto 2021; Summerfield and De Lange 2014).

Finally, the flexibility of our maze tasks provides opportunities to investigate prospective planning across different levels of complexity and experience. Maze structure, the number of branching points, and the distance to downstream obstacles can be varied systematically to test the limits of prospective evaluation. Such experiments could help distinguish alternative planning algorithms, characterize the contributions of learning and prior knowledge, and establish connections between behavioral strategies and their neural implementation. More broadly, our findings suggest that macaques provide a useful model for studying how prospective cognitive abilities generalize to unfamiliar problems, linking questions in systems neuroscience with related challenges in artificial intelligence.

## Materials and methods

### Subjects and apparatus

Three adult male macaque monkeys (*Macaca mulatta*, Monkeys L, M, and H) performed a maze navigation task in their home cages. All experimental procedures conformed to the National Institutes of Health *Guide for the Care and Use of Laboratory Animals* and were approved by the Institutional Animal Care and Use Committee at New York University.

The task was administered on a touchscreen device (Apple iPad Pro, 9.7-inch display, 1024 x 768 pixels, 60 Hz refresh rate, Apple Inc., Cupertino, CA) mounted to a transparent acrylic box (33 cm *×* 48 cm *×* 21 cm) attached to the monkey’s home cage. Monkeys viewed the screen through the transparent panel and interacted with it through an arm hole (13 cm *×* 9 cm) on the front side of the box (Fig. 1A). A custom iOS application developed in Xcode 11.1 controlled the task and recorded behavior. Monkeys received juice reward from a tube mounted above the arm hole. The location of the hole and reward spout were chosen to encourage a stereotyped and comfortable posture, resulting in stable positioning of the head, shoulders, and torso throughout task performance. Head position was 35*±*1 cm from the screen, yielding a visual angle of ∼ 21° (width) *×* 16° (height) for the maze display.

### Maze navigation task

Monkeys performed a maze navigation task (Fig. 1B), in which they moved a token from a starting position on one side of the screen to a goal region on the opposite side while avoiding three immobile obstacles. We refer to obstacles as “walls.” The token, depicted as a banana icon (size, 8% of screen width), could not pass through walls but could be maneuvered above or below them, depending on the size of the vertical gap between each wall and the maze boundaries (indicated by horizontal brick patterns). Each wall had at least one passable gap. The choice of route around one wall (above or below) determined the token’s starting position for approaching subsequent walls. Therefore, the task examined the monkey’s ability to plan multi-segment paths prospectively.

Each trial began with the presentation of a randomly configured maze composed of three walls and a goal region. The token was placed on the side of the screen opposite the goal. For monkey L, its vertical starting position was fixed midway between the upper and lower maze boundaries. For monkeys M and H, the starting vertical position was randomized on each trial, sampled from a uniform distribution spanning 10-90% of the screen height. The goal region was indicated by a vertical green bar extending the full screen height. For monkeys L and M, who performed the task with the right hand, the goal was placed on the left side of the screen. For monkey H, who performed the task with the left hand, it was placed on the right side.

Each wall occupied 50% of the screen height. Their vertical positions were randomized across trials, subject to constraints that defined the specific wall configurations used for training and testing (see below). Horizontally, the walls were spaced to allow ample room for token movement: wall 1 was placed within 30-35% of screen width, wall 2 within 50-55%, and wall 3 within 70-75%. Once the maze appeared, the monkey could move the token toward the goal by touching and swiping. Only swipes initiated on the token were registered as valid; touches elsewhere had no effect. When the token successfully reached the goal, the screen displayed a picture of water droplets (reward cue), and juice was dispensed from the reward tube. If the monkey failed to reach the goal within 30 seconds, the trial was aborted, and no reward was given (*<* 1% of trials). Only successfully completed trials were included in analyses.

Each wall could be configured to allow passage of the token only from above (upper route, U), only from below (lower route, L), or both (B). Consequently, mazes could be classified into 27 distinct configurations (3^3^). Because the vertical wall positions were uniformly sampled, these configurations occurred with unequal prior probabilities. We denote each maze configuration using three-letter codes: for example, ULB indicates that wall 1 allows only the upper route, wall 2 only the lower route, and wall 3 both. Similarly, we use the letters U and L to describe the monkey’s chosen path around each wall.

Prior to the data collection, monkeys were trained in a stepwise fashion to interact with the touchscreen and understand the task structure. Initially, they were rewarded for simply touching the token on a blank screen. Next, they learned to move the token to a goal region in the absence of walls. A single wall was subsequently introduced, and training continued until monkeys could reliably navigate around the wall to reach the goal. This staged training familiarized the monkey with the logic of the task—movement constraints and goal-seeking behavior—without requiring prospective planning. Subsequently, we introduced a second wall, followed shortly by a third wall, which marked the beginning of the full task and the start of data collection. All trials involving three-wall mazes were included in the main analyses. The dataset included 72 sessions across three monkeys (monkey L, 16; M, 30; H, 26), comprising a total of 95,431 trials (monkey L, 34,991; M, 33,028; H: 27,412).

### Control tasks

To test the monkey’s ability to generalize their strategy to new environments, we conducted two control experiments with monkeys M and H. These controls tested generalization over both task-irrelevant and task-relevant changes.

The first control examined generalization across a task-irrelevant feature: wall texture. In the main task, the three walls had distinct brick textures that were presented in a fixed horizontal order (Fig. 1B). In the control task, the order of these textures was randomized, presenting the monkey with novel sequences of wall textures. If the monkey relied on memorized associations between specific visual displays and actions, randomizing textures would disrupt performance. In contrast, if their strategy involved prospective planning based on wall positions, their performance should remain stable despite the novel texture arrangements.

The second control tested generalization to novel maze configurations. For monkeys M and H, we withheld eight of the 27 possible maze configurations during the main task. Monkey H never experienced these eight configurations, while monkey M encountered them only sparsely (1.7% of main task trials). We then introduced these eight configurations in the control sessions. For monkey M, the test configurations were: LUU, LUL, BLU, ULB, BLB, UBB, LBB, and UBU, and for monkey H: UUL, LUL, ULB, BLU, BLB, BBU, BBL, and UBU. These sets included a range of structural patterns, including mirror-image counterparts of familiar mazes, and served to test whether monkeys could generalize their strategy to previously unseen maze configurations. In particular, this control *quantitatively* assessed whether the monkey’s decision around one wall remained appropriately influenced by the configuration of downstream walls, even in novel contexts (see Data Analysis).

Monkeys completed 11 control sessions with randomized wall texture orders (monkey M, 4; H, 7), comprising 12,756 trials in total (monkey M, 4,810; H, 7,946). They also completed 24 sessions with novel maze configurations (monkey M, 10; H, 14), contributing 26,741 trials (monkey M, 11,778; H, 14,963).

### Data analysis

To analyze the monkey’s behavioral strategy, we first identified the monkey’s chosen route around each wall (U or L) based on which gap the monkey moved the token to. If the token entered a gap or touched the wall corner adjacent to the gap, we registered it as the chosen route. In some cases, monkeys initially approached one gap before changing course to the opposite side—typically when realizing that the first gap was impassable. Because these corrections based on visual feedback are not informative about the monkey’s prospective planning, we quantified the monkey’s choice based on the first committed attempt to pass the wall.

We examined the effect of wall positions on the monkey’s choice by plotting the probability of selecting the upper route as a function of each wall’s position (Figs. 1D, 2B, 2D, 4A, S1, S4). For monkey L, we included all trials. For monkeys H and M, we removed a subset of trials to account for the imbalance in the data distribution due to the exclusion of eight wall configurations during the main data collection sessions (see “Control tasks”). For example, excluding the LUU configuration from the main task reduced the frequency of LU wall pairs compared to UL, inducing a spurious correlation between the positions of wall 1 and wall 2, which could artificially inflate correlation in behavior. To correct for this, we randomly sub-sampled trials to equalize the number of U and L trials for each relevant wall when plotting conditional choice probabilities.

To quantify how wall positions influenced the monkey’s selected route around the first wall, we used a logistic regression model:

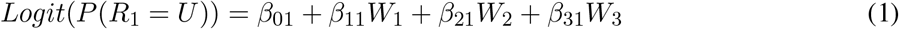

where *P* (*R*_1_ = *U* ) is the probability of choosing the upper route at wall 1, *W*_1_, *W*_2_, and *W*_3_ are the normalized vertical positions of walls 1-3 (ranging from –0.5 to +0.5), and *β*_01_ captures the monkey’s baseline choice bias. *β*_11_, *β*_21_, *β*_31_ are the model coefficients that quantify the sensitivity of the monkey’s choices to the three wall positions. We relaxed trial-balancing constraints for this analysis, as they had minimal influence on the fitted coefficients.

To explore whether a constraining position for wall 2 reduced the influence of wall 3 on the choice around wall 1, we extended the wall-specific model by adding an interaction between wall 3 and a two-state function of wall 2:

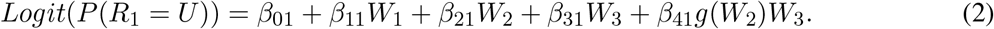

where the two-state function *g*(*W*_2_) was defined as 1 when wall 2 permitted only the upper or lower route (U or L) and 0 when it permitted both routes (B). In this formulation, *β*_41_ quantifies the change in the effect of *W*_3_ when *W*_2_ restricts passage to a single route relative to when both routes are available. We compared this extended model with Eq. 1 using a likelihood-ratio test with one degree of freedom and evaluated the significance of *β*_41_ using a two-sided Wald test. The model was also fit separately to each monkey to assess consistency across subjects.

To test whether first-route choices could be explained by an order-free visual summary of the maze, we devised alternative models. These models discarded the identity and serial position of the individual walls. In the “average-position model” we used the average of the normalized position of the three walls

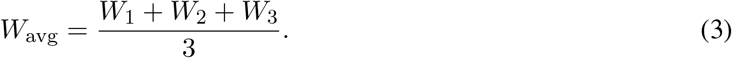

as model predictor. The “largest-gap” model used the difference between the largest upper and lower gaps in the maze as its predictor. For each wall, the corresponding normalized upper- and lower-gap sizes were

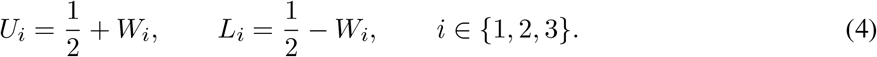

A signed gap score can be defined as the difference between the largest upper gap and the largest lower gap across the three walls:

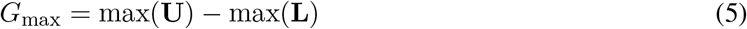

Positive values of *G*_max_ indicate that the largest upper gap was wider than the largest lower gap, whereas negative values indicated the opposite.

To assess whether the influence of downstream walls on first-route choices could be explained by summary measures of upper and lower lane openness, we compared the wall-specific model with the average-position and largest-gap models. We used the same first-route choice observations analyzed with Eq. 1 and performed leave-one-session-out cross-validation separately for each monkey (H, six sessions; L, three; M, six). A session was a single experimental recording period for one monkey. In each fold, each candidate model and an intercept-only null model were fitted by maximum likelihood to all but one session from that monkey and evaluated on the held-out session. Each session served once as the test set. Predictive performance was summarized using McFadden’s pseudo-*R*^2^ calculated from held-out log likelihoods:

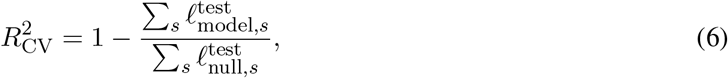

where 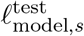 and 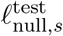 were the log likelihoods of choices in held-out session *s* under the candidate and intercept-only null models, respectively. The sums were taken across sessions within each monkey.

We applied the same procedure to monkey L’s switch-track subset, comprising BUL (n=228) and BLU (n=230) trials. Both configurations occurred in each of three sessions. Model formulas were unchanged, but fitting and evaluation were restricted to BUL and BLU trials for both the candidate and null models. Wall positions were not binned for this comparison.

For the second and third walls, we used similar models but included the monkey’s previous choices as additional predictors. The logistic regression models were:

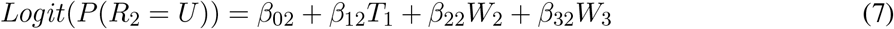

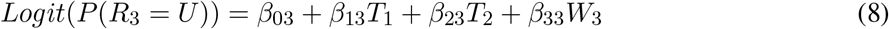

where *T*_*n*_ is a binary variable indicating whether the upper route was taken for wall *n* (1 for U and 0 for L). These models were fit separately for each monkey using maximum likelihood estimation. The goodness of fit was assessed by calculating McFadden’s pseudo-*R*^2^.

The models above assume that the weight of evidence conferred by each wall changes linearly with its position. Alternatively, the weight of evidence may be a non-linear function of wall position. Nonlinearities are plausible because wall positions pose hard boundaries: the token can pass all gaps larger than the token size and fails to pass all smaller gaps. Therefore, the wall position could strongly influence the monkey’s choice when gap sizes are close to the token size, but have a diminished effect for much smaller or larger gap sizes. To test for possible nonlinearities, we applied a sigmoid transformation to wall positions:

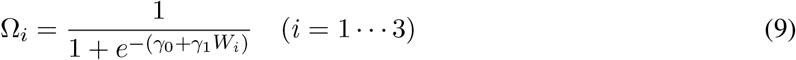

The transformed values, Ω_*i*_, were then used in a modified logistic regression for the monkey’s choice:

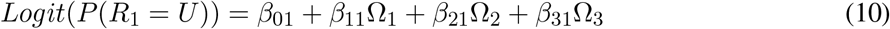

This nonlinear model introduced two additional parameters (*γ*_0_, *γ*_1_). Fits were obtained via maximum likelihood estimation.

To assess how behavior evolved with experience (Fig. 4C-F), we divided the trials of the main task into deciles (2,853-3,502 trials per bin) and computed four behavioral metrics: (1) the probability of choosing the lowest-switch route at the first wall (defined in Results), (2) maze completion time (from the first touch to goal arrival), (3) average swipe speed (pixels/sec), and (4) number of swipes per trial. We modeled changes of these metrics using linear regressions:

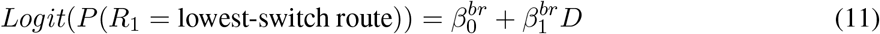

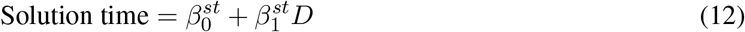

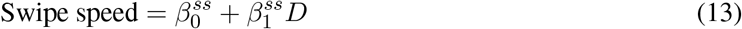

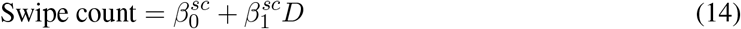

where *D* ∈ *{*0, 1, …, 9*}* indicates the decile. The intercepts (*β*_0_) represent the initial performance, and the slopes (*β*_1_) quantify the rate of change of each metric with accumulated experience.

### Analysis of behavior in multicursal mazes

We analyzed performance in the 6 *×* 6 and 7 *×* 7 mazes for monkeys H and M. In rare cases where a maze structure repeated for a monkey, we retained only its first presentation in the analysis. Completion probability was the proportion of retained trials with a recorded response in which the token reached the goal (Fig. 3B).

To quantify route selection, we analyzed branching points that offered both a goal-reaching route and at least one dead-end route among the available non-backtracking options (Fig. 3C). A non-backtracking option was defined based on movements to adjacent, accessible cells other than the cell from which the monkey had just arrived. A route was classified as goal-reaching if it allowed the token to reach the goal without returning through the current branching point. Otherwise, it was classified as a dead-end route. Because the mazes were designed to have no loops, each eligible branching point had exactly one goal-reaching non-backtracking option. Both successfully completed and unsuccessful trials were included. We analyzed only the first encounter with each physical branching point within a trial and excluded encounters at which the monkey immediately returned to the preceding cell.

To assess whether route selection could be explained by a local goal-direction heuristic, we examined branching points where this heuristic favored a dead end. The heuristic selected the available non-backtracking step forming the smallest angle with the straight-line vector from the branching point to the goal. We calculated angles of available options using the centers of adjacent maze cells in screen coordinates (Fig. 3D). We defined a deceptive dead end as a dead-end branch whose initial step formed a uniquely smaller angle with the goal direction than the other available non-backtracking steps. Branch-point encounters satisfying this condition were classified as deceptive dead-end encounters. Encounters with tied minimum angles were excluded because the heuristic did not specify a unique preferred option. For deceptive dead-end encounters, we measured the probability of selecting a goal-reaching route as a function of dead-end branch depth (Fig. 3E). Depth was defined as the maximum number of maze-cell transitions from the branching point to a terminal cell within the dead-end branch, including the initial transition out of the branching point. When the branch contained additional branching points, we considered terminal cells across all of its subbranches. Dead-end depth quantified the maximum extent of the branch and was used as a proxy for the amount of downstream path structure potentially relevant to identifying the dead end.

For each included branch-point encounter *i*, the node-specific random-choice probability was calculated as *p*_*i*_ = 1*/k*_*i*_, where *k*_*i*_ was the number of available non-backtracking options. The chance level for each analysis was the mean of these probabilities across its included encounters. To test whether goal-reaching choices exceeded chance, we used a one-sided *z*-test based on the sum of observed choices relative to their node-specific random-choice probabilities::

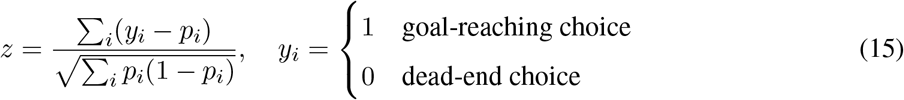

We estimated 95% confidence intervals for completion probabilities, goal-reaching choice probabilities, and choice-minus-chance differences, using a session-cluster bootstrap with 100,000 resamples. Sessions were defined by monkey and date. Within each monkey, sessions were resampled with replacement, retaining all trials and branch point encounters from each session. For each bootstrap resample, we recalculated the relevant probabilities and chance levels, pooling observations across monkeys when applicable. Confidence limits were defined by the 2.5th and 97.5th percentiles of the bootstrap distribution. For the dead-end depth analysis, resampling was performed separately for each depth group.

## Data and code availability

Data and code used in this study will be made publicly available upon official publication.

## Acknowledgements

We thank Lana Sidani and Natasha Recoder for helping with data collection and animal care. We thank Larry Maloney and Elad Schneidman for helpful discussions. This work was supported by the Simons Collaboration on the Global Brain (988247 and GB-Culmination-00002986-02) and Pew Innovation Fund. G.O. was supported by a post-doctoral fellowship from the Charles H. Revson Foundation.

## Supplementary Figures

**Figure S1.**
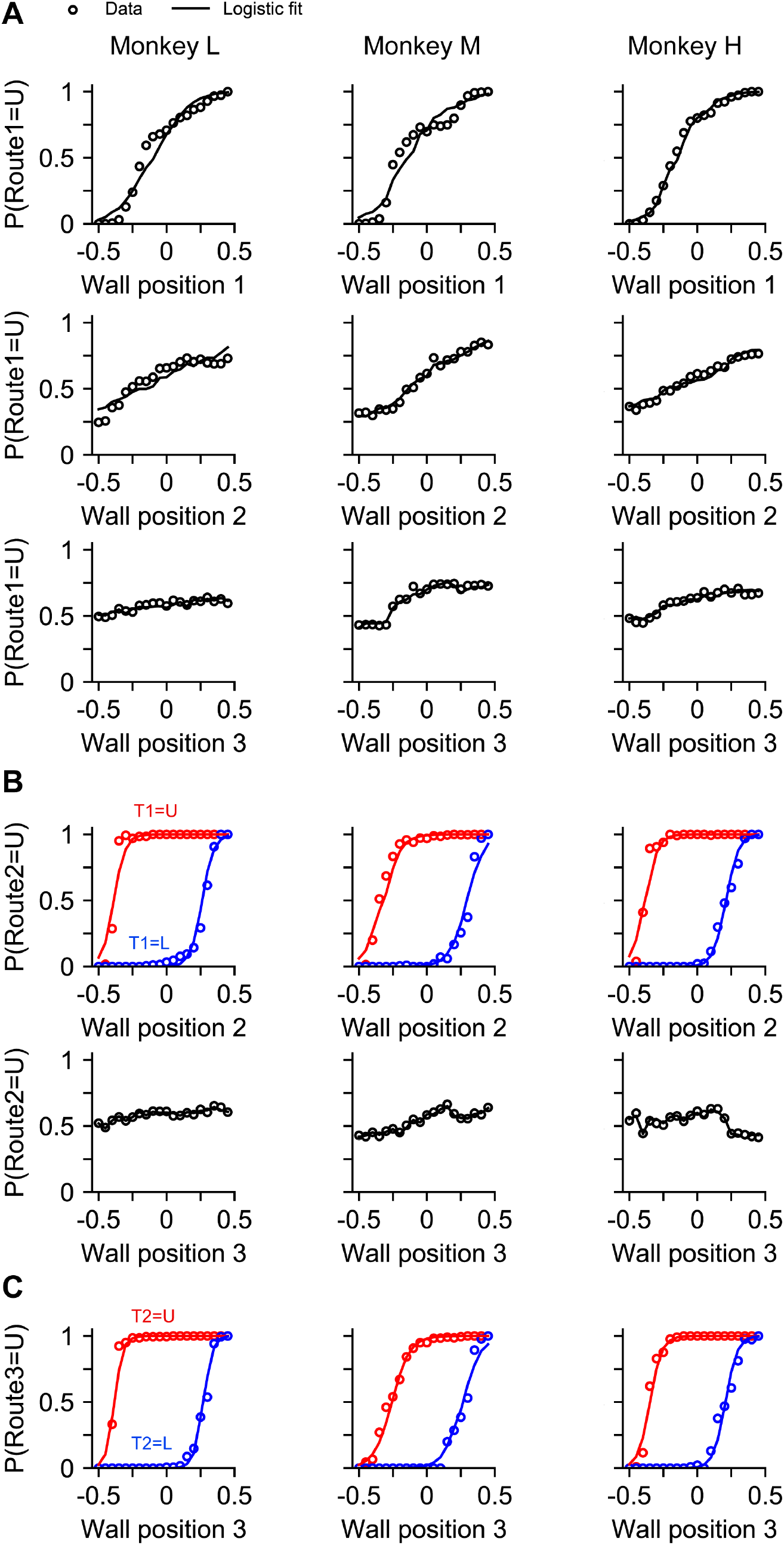
Logistic regression fits for individual monkeys. (A) Choice probabilities of each monkey as a function of wall positions (cf. Figs. 1D, 2B, and 2D for averaged results across subjects). Each column corresponds to one monkey. Open circles show behavioral data, and solid lines indicate logistic model fits (Eqs. 1, 7, and 8). All three wall positions significantly influenced choices at the first wall for all monkeys. **(B-C)** Choices for the second (B) and third (C) walls depended strongly on the routes previously taken around the first and second walls.

**Figure S2.**
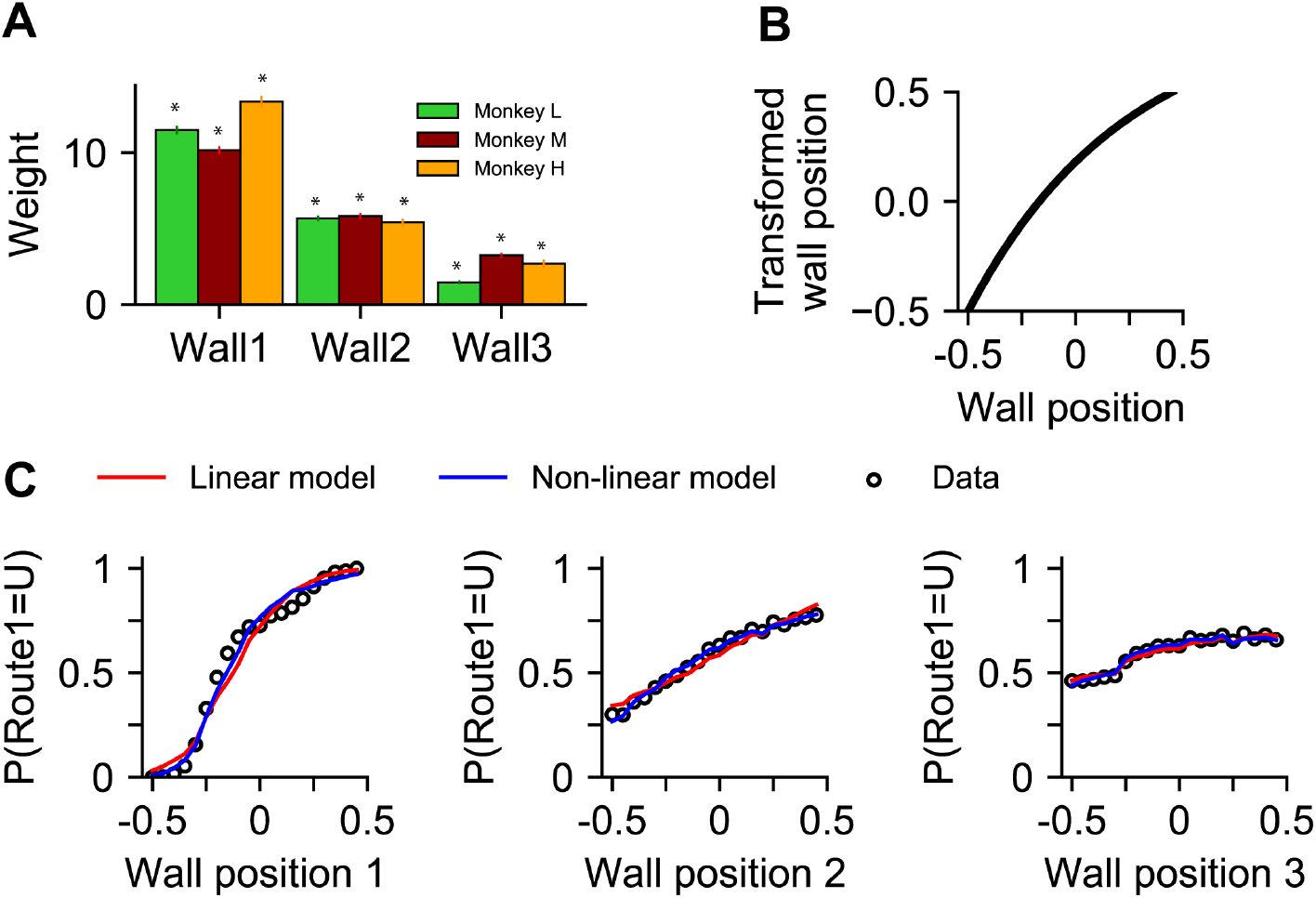
Comparison of linear and non-linear model fits. (A) Logistic regression weights for the linear model (Eq. 1) predicting the route around the first wall as a linear combination of all three wall positions. Model coefficients were comparable across monkeys and significantly greater than zero, indicating that all walls influenced monkeys’ choices. Asterisks indicate p < 0.01. Error bars represent SEM. **(B)** To account for potential nonlinearities—e.g., potential saturating effects for very small or very large wall gaps—we applied a sigmoid transformation (Eq. 10). However, the transformed wall positions that best explained the choices showed no strong non-linear trend. For visualization, transformed wall positions are rescaled to range from –0.5 to 0.5. **(C)** The linear and nonlinear model fits are nearly identical, indicating that the added non-linearity did not substantially improve the model’s explanatory power.

**Figure S3.**
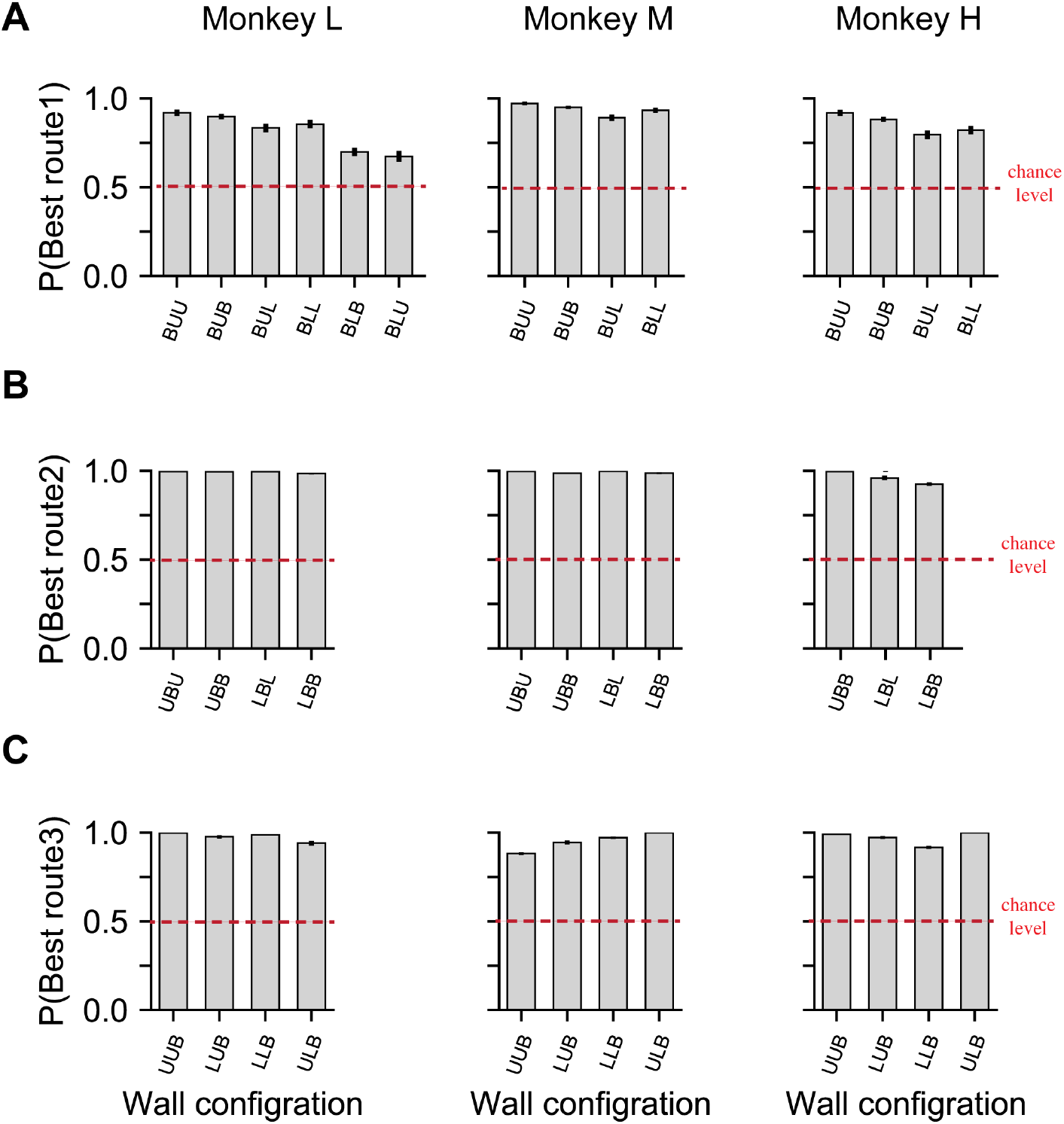
Probability of choosing the lowest-switch routes for individual monkeys. Each column corresponds to one monkey. Some wall configurations were withheld in the main experiments for monkeys M and H to be used in control experiments for testing generalization (see Methods). Monkey L’s results are identical to those shown in Figs. 1E, 2C, and 2E.

**Figure S4.**
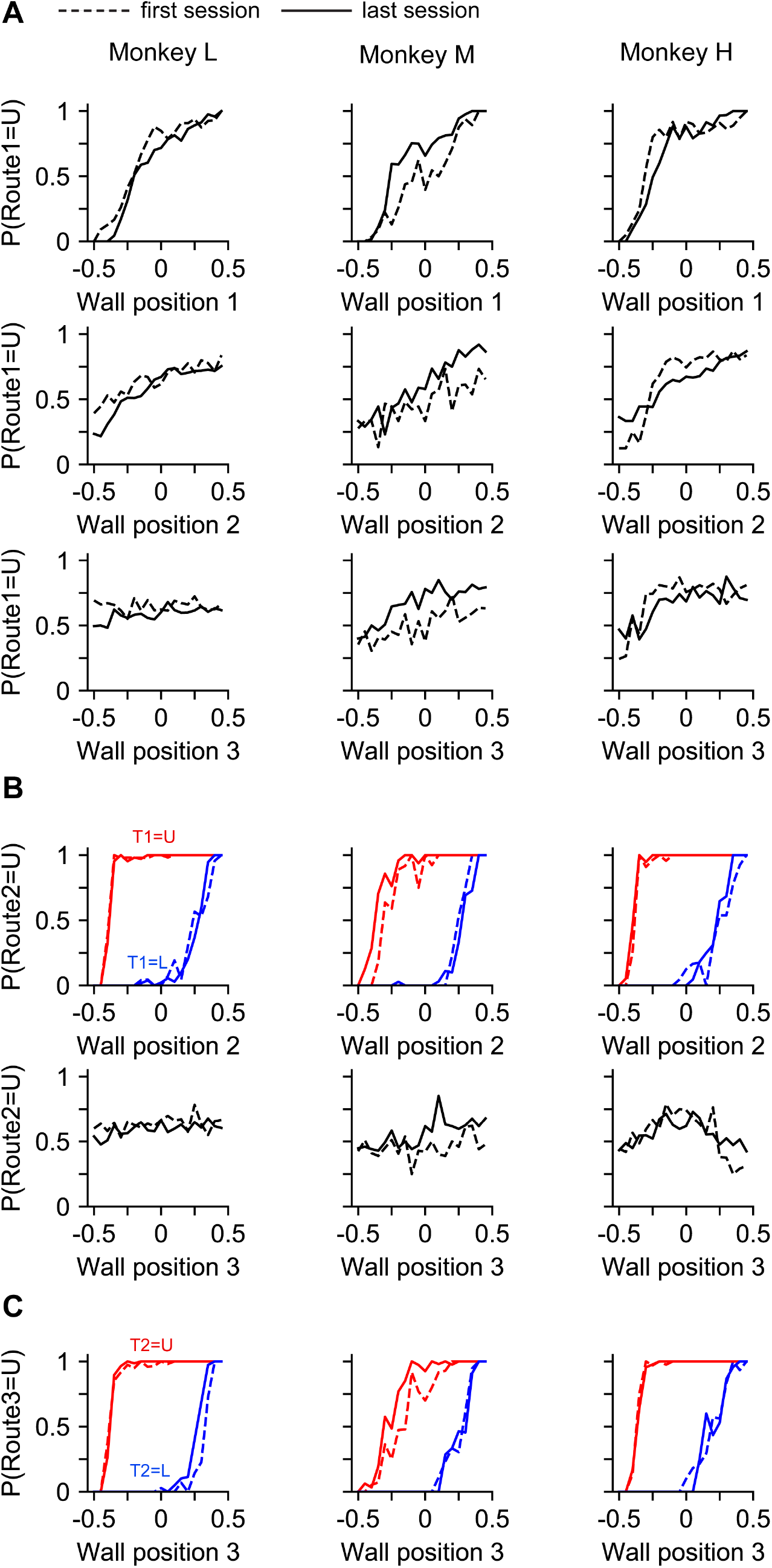
Individual monkey’s behavior was comparable between the first and last sessions of the main experiment. Each column corresponds to one monkey. Logistic regression fits from the first and last sessions for each monkey (cf. Fig. 4A for averaged data). While some differences are visible (e.g., in monkey M), they primarily reflected a shift in the baseline choice bias between sessions. The slopes of the psychometric functions remained largely stable, indicating that the effect of wall positions on monkeys’ choices was consistent over time.

## Notes

### Competing Interest Statement

The authors have declared no competing interest.

